# Empirical and Computational Evaluation of Hemolysis in a Microfluidic Extracorporeal Membrane Oxygenator (ECMO) Prototype

**DOI:** 10.1101/2020.06.15.152686

**Authors:** Nayeem Imtiaz, Matthew Poskus, William A. Stoddard, Thomas R. Gaborski, Steven W. Day

## Abstract

Microfluidic devices promise to overcome the limitations of conventional hemodialysis and oxygenation technologies by incorporating novel membranes with ultrahigh permeability into portable devices with low blood volume. However, the characteristically small dimensions of these devices contribute to both non-physiologic shear that could damage blood components and laminar flow that inhibits transport. While many studies have applied empirical and computational models of hemolysis to medical devices, such as valves and blood pumps, little is known about blood damage in the microfluidic flow regime. In this study, four design variants of a microfluidic membrane-based oxygenator and two controls (positive and negative) are introduced and modeled using a Computational Fluid Dynamics (CFD) model to predict hemolysis. The simulations were performed in ANSYS Fluent for nine shear stress-based Power Law hemolysis model variants. Empirical testing of the devices in a recirculating loop revealed levels of hemolysis significantly lower (3 ppm hemolysis for pump, tube, and device combined) than the hemolysis ranges (>10 ppm) observed in conventional oxygenators. We found that most of the nine tested hemolysis models overpredict (5× to 10×) hemolysis compared to empirical experiments. However, two models demonstrated higher predictive accuracy for hemolysis values in devices characterized by low shear conditions, while another set of three models exhibited better performance for devices operating under higher shear conditions. Our study highlights the limitations of combining hemolysis models with computational fluid dynamics models for a priori in silico device-induced hemolysis. Nevertheless, with a judicious selection of hemolysis models based on the shear ranges of the test device, we propose that computational modeling can complement empirical testing in the development of novel micro-dialyzers or oxygenators, allowing for a more efficient iterative design process. Furthermore, the low device-induced hemolysis (< 2 ppm) measured in our study at physiologically relevant flow rates is promising for the future development of microfluidic dialyzers and oxygenators.

## 3. Introduction

The prevalence of chronic lung diseases, such as Chronic Obstructive Pulmonary Disease (COPD), and unpredictable and potentially overwhelming outbreaks of acute infectious illnesses, such as swine flu and COVID-19, emphasizes the need for improved treatments for respiratory insufficiency and respiratory failure.[1]–[4] Currently, the standard of care is mechanical ventilation, which involves invasive procedures that carry significant risks, including barotrauma, ventilator-associated pneumonia, and other infections.[5], [6] Furthermore, the complications linked to administering mechanical ventilation to manage novel pandemic illnesses such as COVID-19 have escalated the significance of Extracorporeal Membrane Oxygenation (ECMO), a treatment method that uses an external circuit to circulate blood for gas exchange in an artificial lung, enabling the possibility of reducing or, in rare cases, circumventing the requirement for ventilator support.[7]–[10]

Microfluidic-based devices with ultra-high permeability membranes could revolutionize the Extracorporeal membrane oxygenation (ECMO) process by reducing blood volume, blood contacting surface area, and overall device size. Reduced blood volume is helpful to all patients and critical for smaller patients, such as neonates, as the priming volume of conventional medical devices sometimes exceeds the total blood volume of a neonate.[11]–[14] Additionally, the associated decrease in membrane surface area and blood-contacting artificial materials reduces complications and blood damage.[15]–[17] Conventional oxygenators are typically a tube-in-tube configuration,[18], [19] whereas microfluidic devices are often arranged in stacked planar plates, an architecture that lends itself to a family of recently developed ultra-high permeability membranes.[15], [16], [20], [21] At physiological flow rates and at flow rates observed in conventional ECMO devices, the small dimensions and sharp edges of microdevices may result in shear sufficient to induce hemolysis (rupture of red blood cells) and other bleeding disorders.[22]–[24] While many studies have applied empirical and computational models of hemolysis to medical devices, such as valves and blood pumps, little is known about blood damage in the microfluidic flow regime.[25]–[27]

Acute hemolysis is rare in current ECMO treatments, but effects can aggregate with frequent or continuous treatment. The devices analyzed in this study are microfluidic [28] ECMO devices with micrometer-to-millimeter scale channel widths and flow rates of 100s of ml min^-1^, which are substantially slower than conventional ECMOs [29], yet are orders of magnitude larger and higher flow than lab on chip microfluidics.[28], [30] Devices at this scale represent a “middle ground” where the design community has limited experience predicting the device-induced/ hemolysis (Fig. 1). Furthermore, common models to predict hemolysis were empirically generated in Couette flow shearing devices and may not be suitable for predicting flow in regimes found in microfluidic ECMOs.

**Fig. 1.**
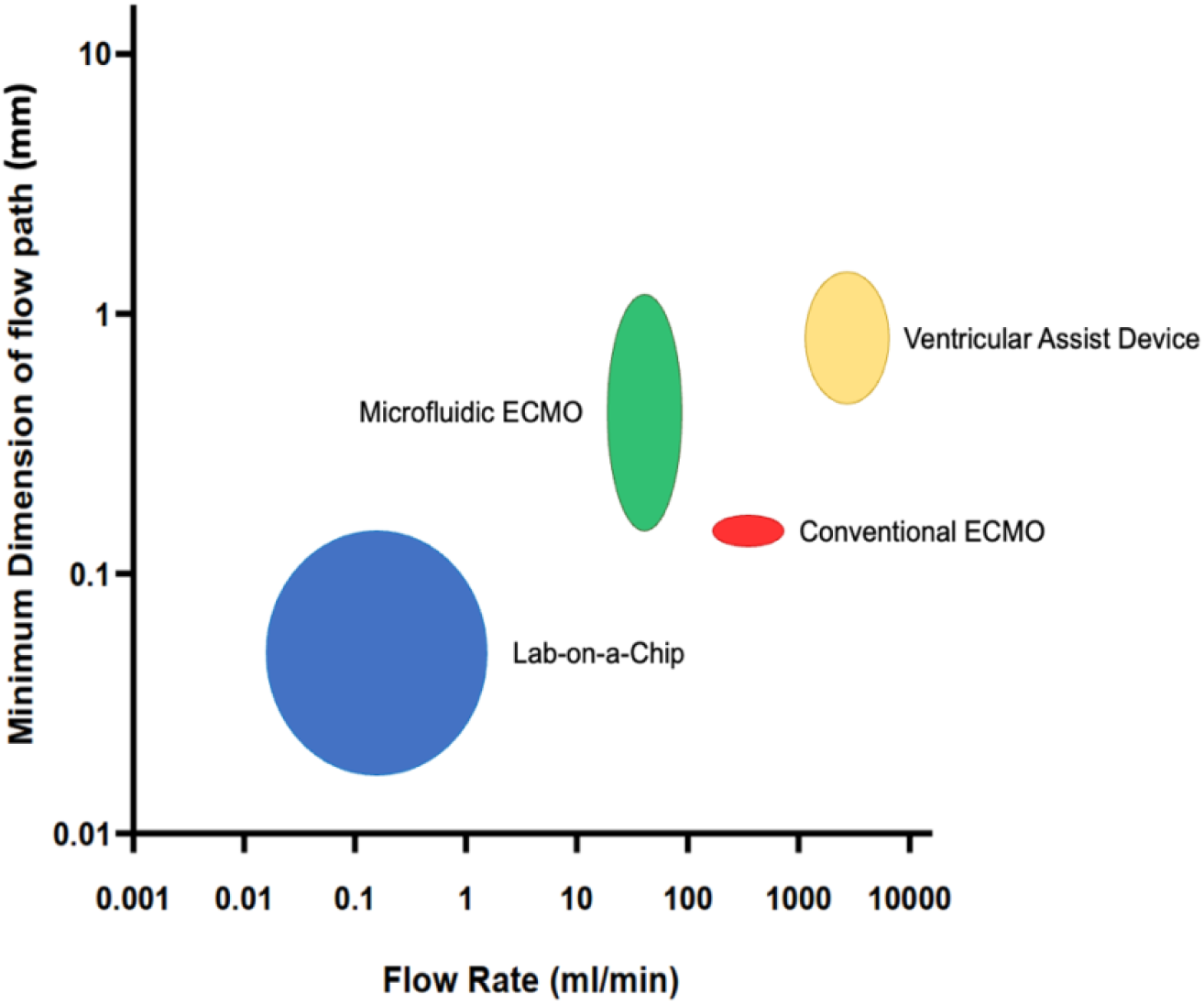
Comparison of different types of blood contacting medical devices (i.e., Lab-on-a-Chip [28], [30], [31], Microfluidic ECMO[17], [32]–[34], Conventional ECMO[29], [35], [36], and Ventricular Assist Device[37]–[40]) in terms of minimum dimension of blood flow path and flow rate through the channel of minimum dimension. The height and width of the circle/oval represents the minimum dimension and flow rate ranges respectively.

Computational modeling is commonly employed during the design process of blood-contacting medical devices.[41]–[43] Hemolysis models are frequently integrated directly into the framework of a computational fluid dynamic (CFD) solver [44], [45], and these methods guide the design process for devices such as microfluidics[46], pumps, valves, and catheters.[47]–[49] A range of hemolysis models exist that vary in complexity, but most predict hemolysis based on empirically determined functions of shear stress *τ*_*s*_ and exposure time *t*, both of which are calculated by CFD. [26], [50] Two methods/classes of implementing hemolysis models are commonly used: 1) Power Law models, where the damage is a result of the product of time and shear to empirically determined constants, and 2) Time History models, which are variants of the Power Law models that consider the fact that RBCs that have previously experienced shear will release less hemoglobin upon subsequent exposure to the same shear.[51] Either of these models is often used to predict the effect of an incremental design variation.[42], [48], [52] It is unclear whether existing hemolysis models for the conventional fluidic devices are suitable for predicting hemolysis for the development of micro-oxygenators and micro-dialyzers, as these devices may have greater or non-uniform shear stress, exposure time, and flow characteristics.

Solute transport within these devices is limited by the transport rate within the laminar fluid rather than the membrane permeability itself. A solute-depleted fluid layer adjacent to the membrane will result in a reduced concentration gradient across the membrane, limiting membrane flux and resulting in insufficient filtration efficiency.[15], [20] Geometrical protrusions have been proposed as means to improve mixing within microfluidic channels (Fig. 2B). [53], [54] Any component of velocity that is perpendicular to the surface acts to move high-concentration fluid into the boundary layer, thus replenishing the depleted boundary layer through advection. Herringbone mixers are chevron-shaped steps in the device that induce countercurrent vortices in the flow.[55]–[57] The staggered herringbone mixer is a well-characterized mixing element that has been demonstrated to improve mixing for a wide range of flow regimes (1 < Re < 100). [55], [58] However, the extent to which these flow disturbances caused by herringbone mixers contribute to flow-induced hemolysis has not yet been studied.

**Fig. 2.**
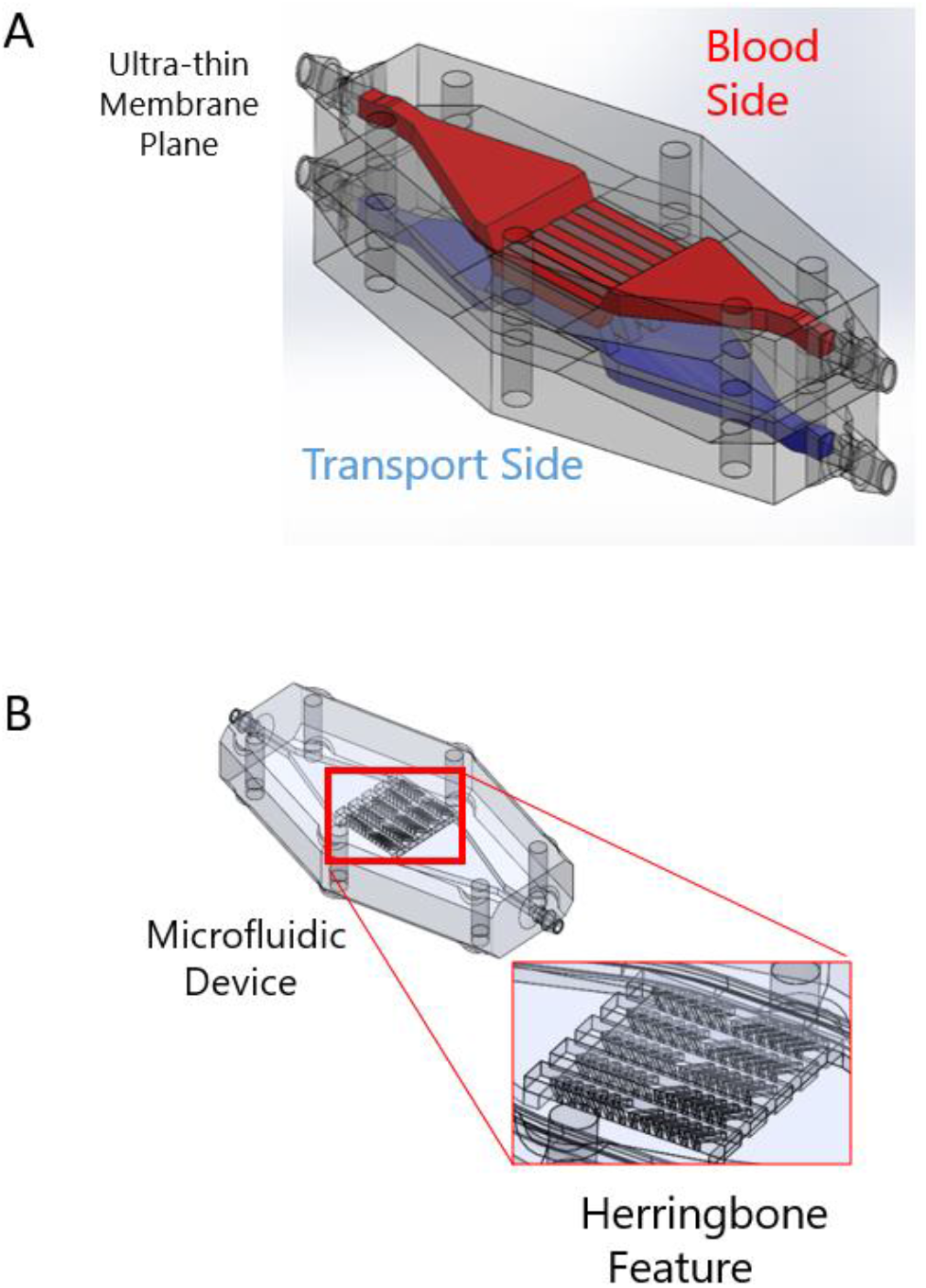
(A) Microfluidic dialyzer with ultra-thin highly permeable membrane, (B) Blood side of the dialyzer with herringbone mixers.

Our study uses a computational model to predict hemolysis in a microfluidic ECMO. Advanced 3D printing techniques were employed to fabricate test device prototypes.[59], [60] In this study, four geometric variants of a prototype microfluidic ECMO were investigated both computationally and empirically to assess 1) the computational model’s ability to predict hemolysis, 2) the comparison of empirical hemolysis with different prototype devices.

## 4. Materials and Methods

### Computational Model Set Up Geometrical model

The geometries of a 5-channel oxygenator device with 1 baseline and 3 variations were created to mirror the experimental setups and test the computational blood damage model against experimental results. The geometry of the baseline configuration is seen in Fig 3. In addition, a geometry was created, simulating the positive control (high shear) case consisting of a 30 mm long, 0.4 mm diameter tube segment between 2 segments of inlet and outlet tube at the 1.6 mm diameter used for the other devices’ ports.

**Fig. 3.**
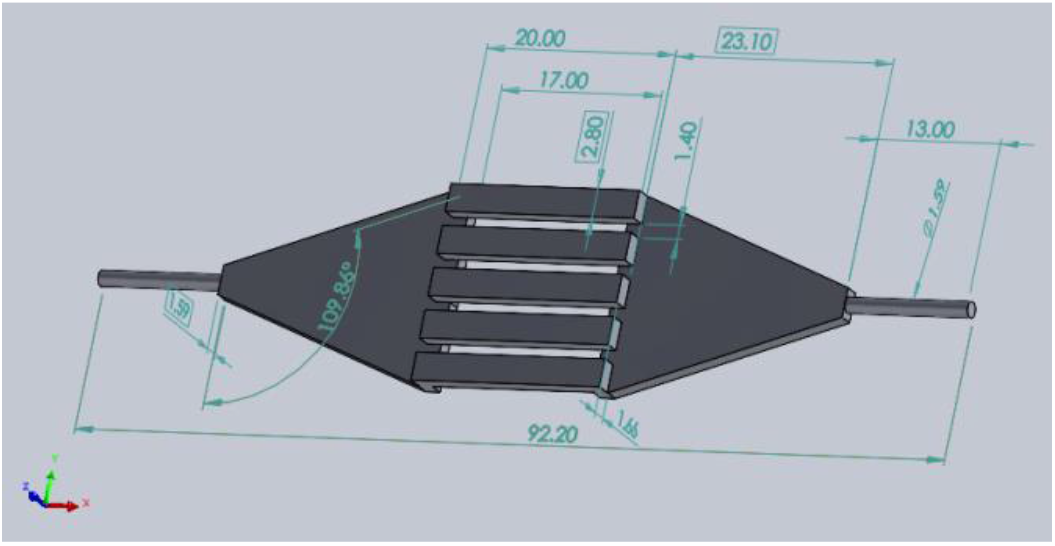
Baseline geometry. Dimensions in mm.

### Numerical Mesh

For each geometry, the solution was resolved for three mesh sizes: 80, 113, and 160 *μm* CutCell (hexahedral) elements (fine, medium, and coarse respectively). Mesh refinement regions proportional to the base mesh size were specified for areas with high-velocity gradients to improve solution accuracy and convergence. (Fig. 4) The shallow channel (250 *μ*m) of one of the test devices (the reduced gap device) required an element size that is even more refined (57 *μ*m) to resolve flow and reach a converged solution. Likewise, the smaller scale of the positive control required the size of the elements to be reduced to 15 *μm*. Each mesh contained approximately 24M hexahedral computational cells at the finest resolution. The average cell orthogonal quality was 0.992.

**Fig. 4.**
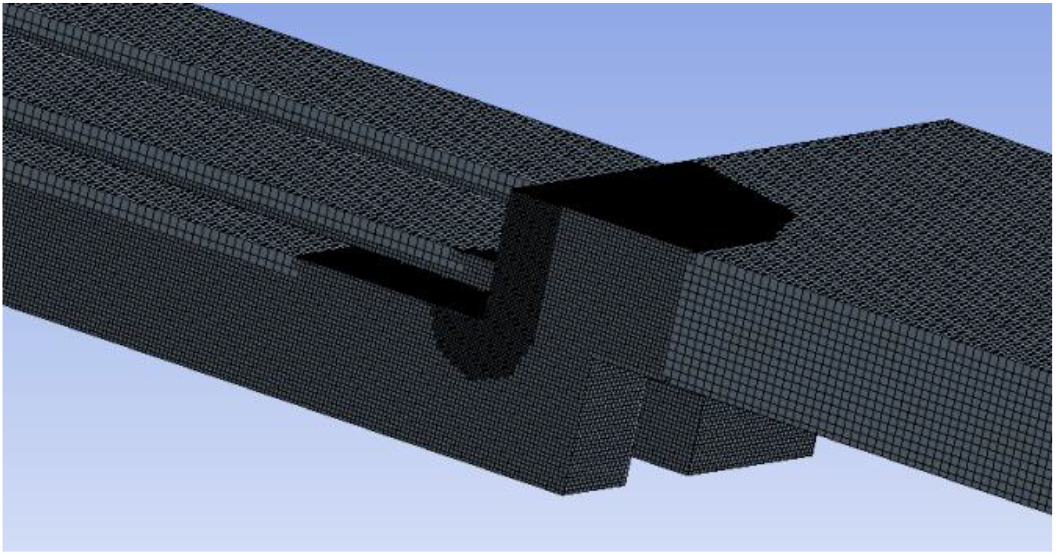
Detail from medium level mesh on baseline geometry.

A mesh convergence study was performed using Richardson Extrapolation to assess mesh discretization error.[61]–[64] For the baseline configuration, the high convergence order suggests that grid-induced error is less than 5%. The results of the mesh convergence study can be seen in Fig. 5. Given the order of magnitude of the differences in hemolysis, 5% grid-induced error is considered acceptable.

**Fig. 5.**
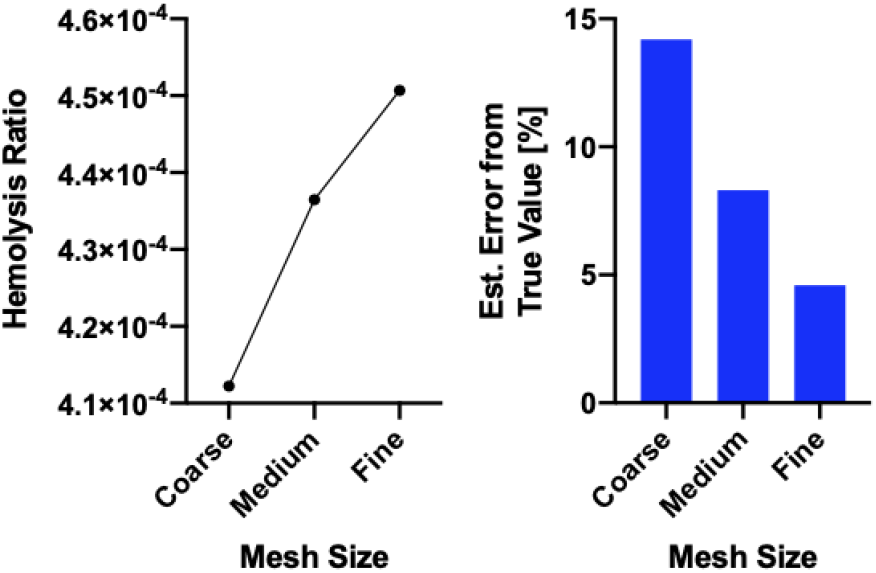
Error calculated using Richardson extrapolation for baseline case.

### Computational Algorithm

A computational model of hemolysis using the native fluid flow solver of ANSYS Fluent and additional User Defined Functions (UDFs) was used. The simulation workflow for the hemolysis is shown in Fig. 6. The core CFD algorithm solved for the fluid dynamics of the system, i.e., blood [65], [66], and the UDF solved for hemoglobin released and free hemoglobin. Cell-free hemoglobin (CFH) was defined as a species. A UDF was implemented to model hemolysis by using a source term to create CFH as a function of shear stress and exposure time, as described in greater detail later.

**Fig. 6.**
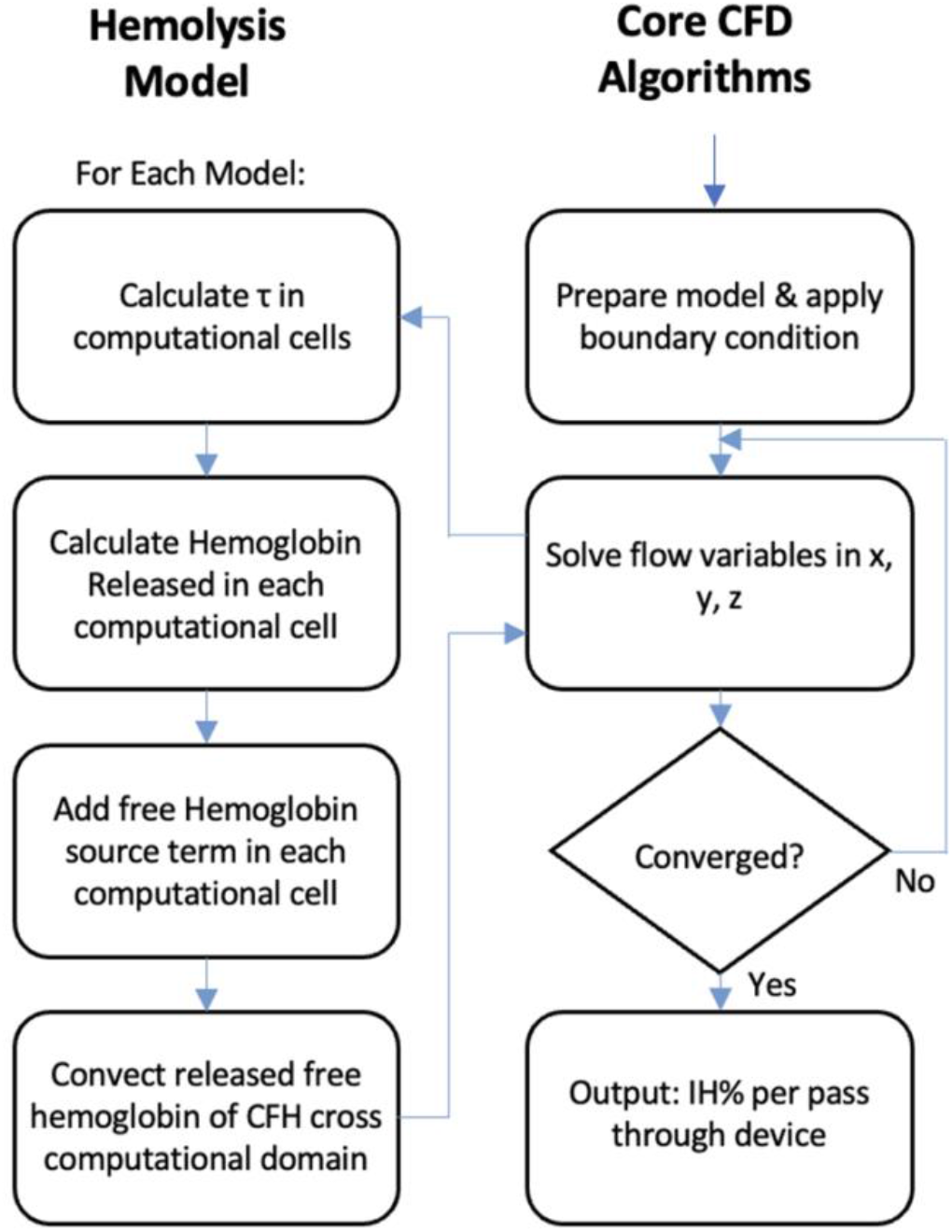
Flow chart of the hemolysis incorporated via UDFs. Fluent solves for fluid flow using the core algorithms (illustrated in the right column). Hemoglobin is treated as a species with a source term that allows for the generation of hemolysis from shear stresses.

A parabolic velocity profile (corresponding to either 100 ml min^-1^ or 10 ml min^-1^) was specified at inlets to avoid non-physical wall shear stress. CFH at the inlet of the device was set to zero. Single-pass damage was quantified by the mass-averaged CFH concentration at the outlet.

The solutions from the simulations were attained at steady-state using a blended first- and second-order upwind scheme (blending factor of 0.75). Convergence criterion of 0.5 × 10^−6^was specified for hemolysis. Flow and hemolysis equations were solved simultaneously on a four-node cluster (4 cores, 32GB memory per node). The computational time to obtain the solution was approximately 24 hours at the finest mesh resolution.

### Hemolysis Model

For each iteration, scalar shear stress was calculated in each computational element. The concentration of a species, in this case hemoglobin, is a scalar value, for which the concentration can be solved at every computational cell in the domain according to (Eqn. 1), where to *ϕ*_*k*_ is the transported scalar quantity. ^44^

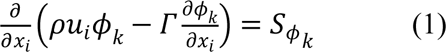

Here, *ρ* is the fluid density, *u*_*i*_ is the velocity of flow along the axial direction, *ϕ*_*k*_ is the transported scalar quantity, and Γ is the diffusion coefficient. The first term *ρu*_*i*_*ϕ*_*k*_ represents the advective term and the 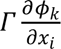 term represents convection. The term on the RHS is the source term, which models the release of free hemoglobin from within cells to the plasma. The source term for each Power Law model and Time History model was defined by the governing Eqs. 2 and 3 respectively. A total of 9 Power Law (PL) models and 9 Time History (TH) were studied.

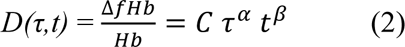

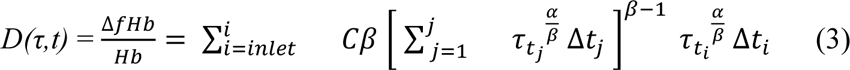

For both Power Law and Time History model three sets of previously published empirical constants, C, *α*, and *β*, were considered in our study. These were published Giersiepen, Heuser/Optiz, and Zhang.[26], [67]–[69] Lastly three different methods of computing the scalar shear stress based on the viscous stress tensor reported by Fluent were employed. Two out of the three methods are based on the second stress invariant (*τ*_*vm*_ and *τ_p_*) and one is based on the viscous stress tensor (*τ*_b_). *τ*_p_ is the square root of the absolute value of the second stress invariant of the deviatoric stress tensor.[70] *τ*_*vm*_is the von mises stress of this stress tensor, or simply √3 ∗*τ*_*p*_. *τ*_b_ is computed from the off-diagonal components of this stress tensor *τ_b_* as shown in Bludszuweit et al.[71] A total of eighteen damage models were studied (PL & TH, 2 * 3 parameter sets * 3 scalar stress for each = 18) (Table 1). The values of the constants C, *α*, and *β* are listed in supplementary table S4. The hemolysis per pass through the device was calculated using Eq. 4 as the average index of hemolysis percentage (IH%) at the device outlet. Ht denotes the hematocrit level in whole blood.

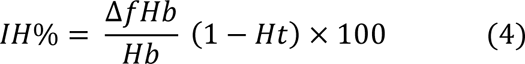

**Table 1:** Hemolysis model parameters.

| Parameter Set | Shear Stress Model |  |  |
| --- | --- | --- | --- |
| | $\tau_{vm}$ | $\tau_b$ | $\tau_p$ |
| Giersiepen | PL-1, TH-1 | PL-2, TH-2 | PL-3, TH-3 |
| Heuser/Optiz | PL-4, TH-4 | PL-5, TH-5 | PL-6, TH-6 |
| Zhang | PL-7, TH-7 | PL-8, TH-8 | PL-9, TH-9 |

### Empirical: Hemolysis

We measured hemolysis in four geometrical variants of a microfluidic oxygenator, as well as positive and negative controls, to explore the model’s predictive power of both absolute magnitude and sensitivity to geometry variations (Fig. 7. Table 2). All the devices contain an inlet manifold which transitions from circular ports to rectangular channels and the rectangular channels subsequently transition into an outlet manifold with circular ports for the outlet. The four geometric variations display variations in both average and maximum shear rates (Table S3), suggesting the possibility of distinct hemolytic potentials for each device. The devices were fabricated by stereolithography (Form 3 SLA Printer, Biomed Amber Resin, Formlabs). Induced hemolysis was measured using a recirculating circuit driven by a custom syringe pump (Fig. 8A, 8B). Empirical experiment setups (test circuits) were constructed of 60 ml syringes (Qosina) attached to a custom-built syringe pump, polycarbonate fittings (VWR), silicone tubing (MasterFlex), microfluidic devices, and blood-bags (100 ml, 2 port blood bag, Qosina). A device-free circuit (1.6 mm ID tubing) was included as a negative control and to establish a suitable positive control, a tubing with a 0.4 mm ID was chosen, as it generates the highest shear levels among all tested devices while remaining within the observed range of shear experienced by blood-contacting devices.[24]

**Fig. 7.**
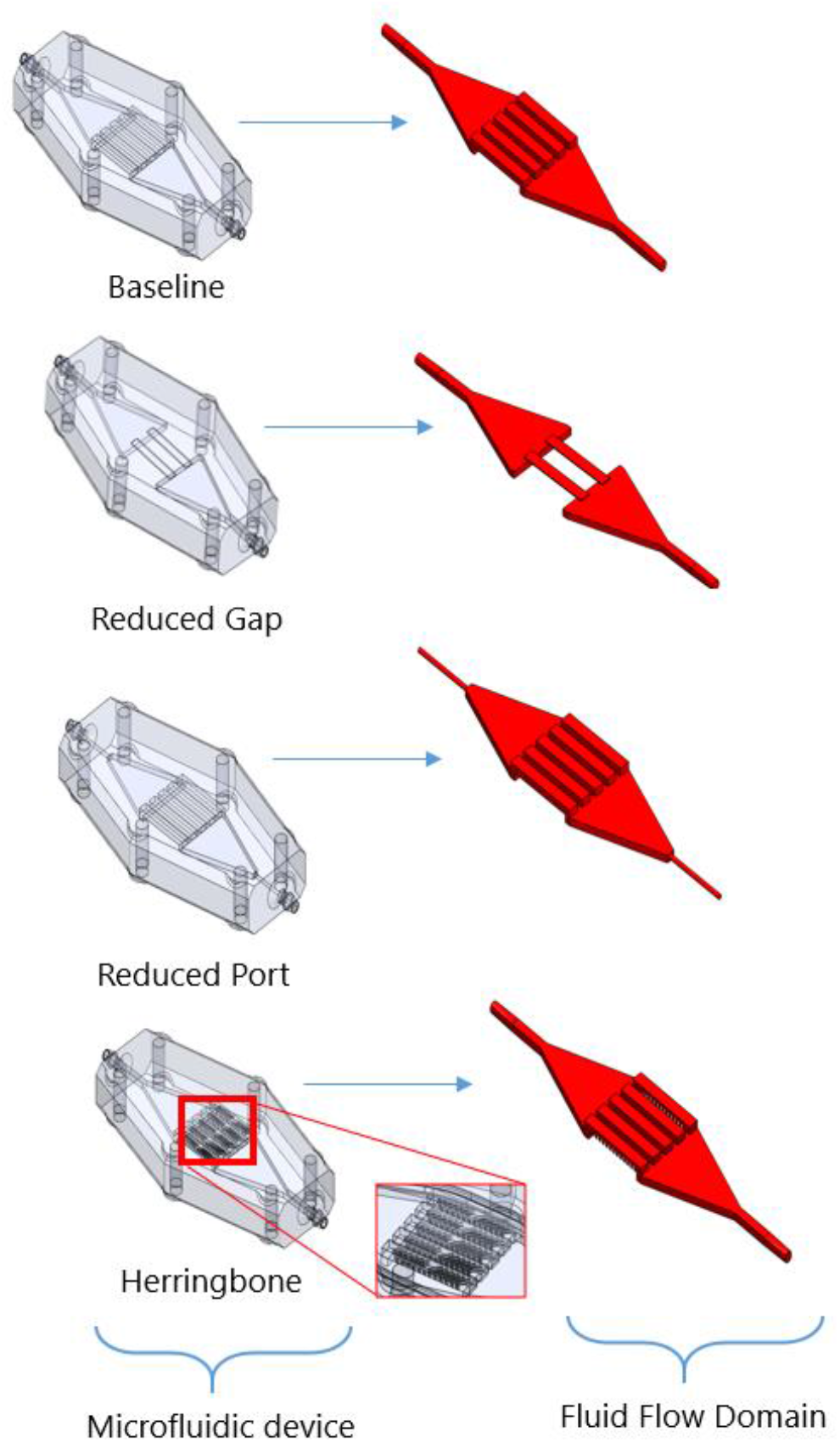
Device variants and their corresponding flow domains. *Baseline:* five open channels, *Reduced Gap:* fewer channels with smallest height to increase shear, *Reduced Port:* small inlet and outlet ports, *Herringbone:* channels with herringbone features to enhance mixing; herringbone height is half of the channel height.

**Fig. 8.**
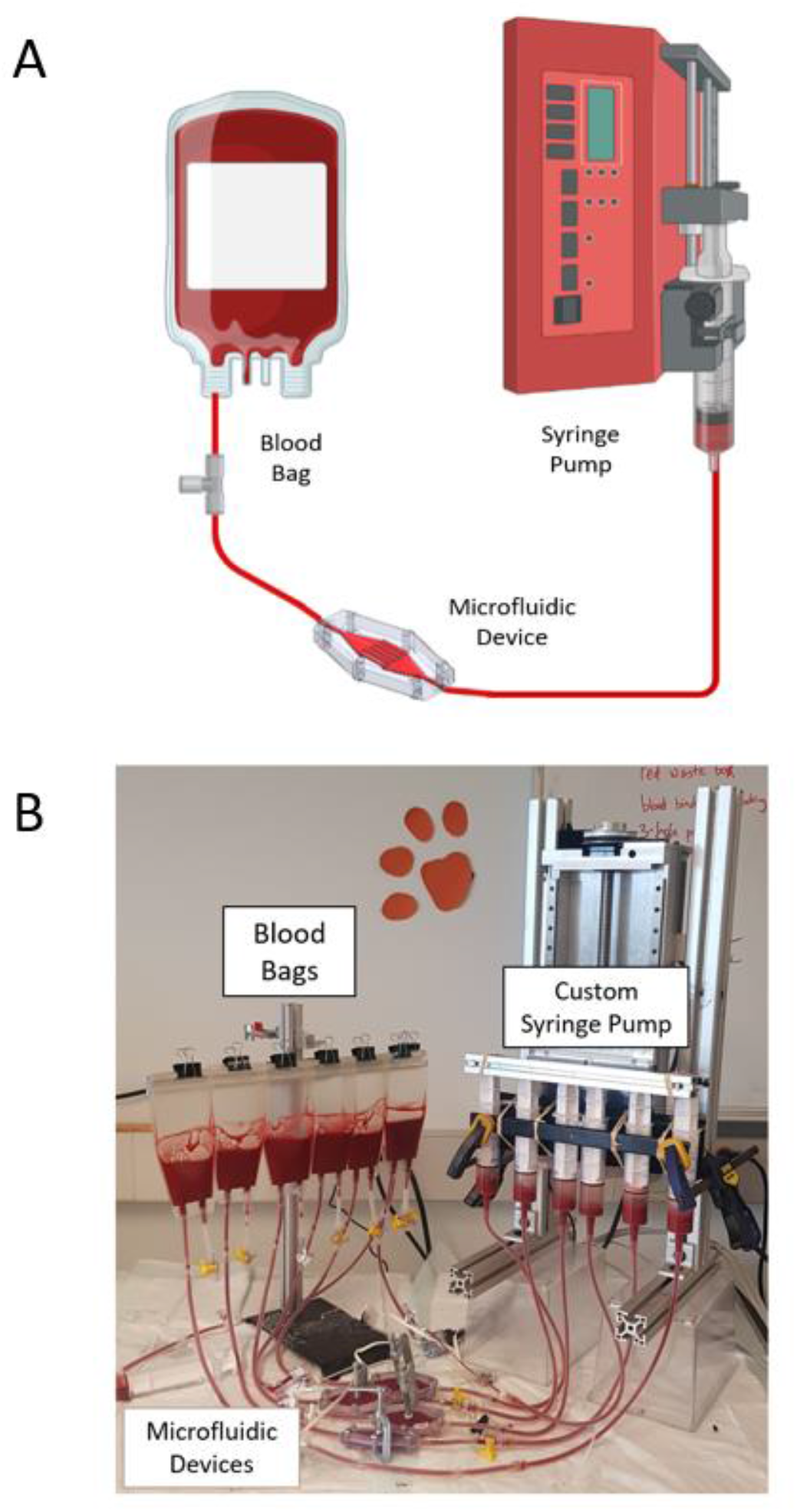
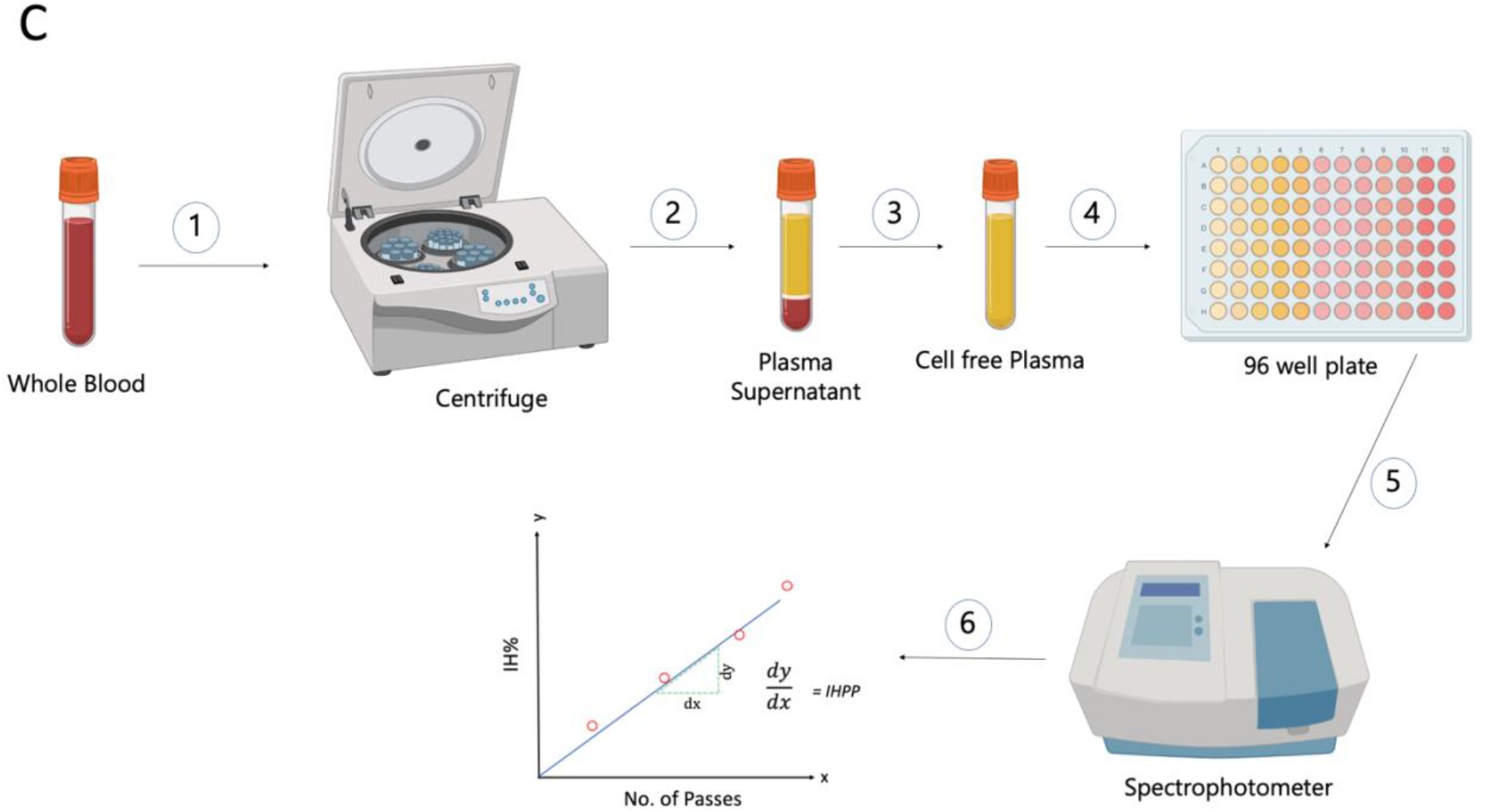
(A) Schematic diagram of the experimental setup. (B) Test circuit with Custom Syringe Pump, Microfluidic Devices, and Blood Bags. (C) Procedure to obtain IHPP from the blood samples.

**Table 2.** Geometrical features of microfluidic devices. The difference from the Baseline device is bolded.

|  | Baseline | Reduced Port | Reduced Gap | Herringbone | Positive Control | Negative Control |
| --- | --- | --- | --- | --- | --- | --- |
| # of Channels | 5 | 5 | 2 | 5 | — | — |
| Channel Width (mm) | 1.6 | 1.6 | 1.6 | 1.6 | — | — |
| Channel Height (mm) | 2.8 | 2.8 | **0.25** | 2.8 | — | — |
| Channel Length (mm) | 20 | 20 | 20 | 20 | 30 | 50 |
| Port ID (device)/<br>Tube ID (control) | 1.6 | **1** | 1.6 | 1.6 | 0.4 | 1.6 |
| # of Herringbones per channel | 0 | 0 | 0 | **12** | — | — |

Citrated bovine calf whole blood was acquired through venipuncture (Lampire Biological Laboratories, PA) from a single donor animal, shipped overnight in a refrigerated container, and used within 12 hours of receipt. Blood was filtered through a 250-micron pore mesh (McMaster Carr). Before each experiment, the circuits were filled with 1X Phosphate Buffered Saline (PBS) for priming and circulated for 20 minutes according to standardized hemolysis testing protocols.[52] The PBS was subsequently removed, and 60 mL of blood was immediately infused into the circuits via the syringe pump. The syringes were programmed to pass blood 650 times through each device at a flow rate of 100 ml min^-1^ for both aspiration and infusion. After every 100 strokes of the syringe, a blood sample of 2 ml was aspirated from each circuit. The samples were centrifuged at 13,000g for 10 minutes and the separated plasma was isolated and centrifuged again to remove any remaining cells. CFH in the plasma was measured according to the Cripps method,[72] which is a well-established method for measuring hemolysis that uses light absorbance values at 560, 576, and 593 nm wavelengths by a spectrophotometer (Spectramax iD3, Molecular Devices). The index of hemolysis (IH) was computed by accounting for the hematocrit. The number of passes was calculated with consideration of circuit volume, flow rate, and total number of syringe strokes. 4 to 7 biological replicates were collected for each device from distinct donor animals. The mean IH was calculated for each device incorporating hemolysis values obtained from all the days of experiments. The standard mean of error was also calculated for each mean IH. A procedure workflow for getting IHPP from blood samples has been shown in Fig. 8C.

### Data Analysis

The slope of a linear regression between IH% and the number of passes through the device or control (IH% pass^-1^, written as IHPP, index of hemolysis per pass) was used as the metric for damage. IHPP from each of the eighteen variations of the hemolysis model were compared to each other and to corresponding empirical data. The cross-device hemolysis comparison for the empirical experiments was done using the one-way analysis of variance (ANOVA), where p < 0.05 indicates that two sets of data are distinct. The computational and empirical data for the devices were compared by one-sample t-test, where a p < 0.05 indicates that the computational datum is distinct from the empirical data.

## 5. Results

### Simulation: Hemolysis

Simulation results of the 18 different hemolysis models for baseline geometry showed little difference between power law and time-history models (Fig. 9A). Being so, only the power law models are shown in Fig. 9B and 9C and hereafter. The magnitude of hemolysis predicted by the various computational models for a single geometry varies drastically (Fig. 9B, 9C), with an overall range of two orders of magnitude difference between Model 3 and 5 (Fig. 9B, 9C). The strongest effect is a result of the parameters *C*, *α*, *and β* derived from the literature, with Giersiepen (Models 1-3) predicting the highest hemolysis and Heuser-Opitz (Models 4-6) the lowest. The means of calculating shear has a weaker effect on the prediction with *τ*_*p*_ shear (Models 3, 6, and 9); consistently predicting 4 times greater hemolysis per pass than the *τ*_*b*_ shear (Models 2, 5, and 8). All models are consistent in predicting the relative changes between devices, namely the herringbones increase hemolysis per pass through the device <10% and all models predict the small port and reduced channel gap devices (high shear) induce a 10 to 20-fold increase in hemolysis compared to the featureless device (baseline). The right-most group in figure 6B depicts the hemolysis values found from the empirical experiments, which will be described in the *Computational vs. Empirical Hemolysis* section. The negative control showed the lowest level of hemolysis in all models when compared to the devices. The positive control had orders of magnitude higher hemolysis compared to the other devices.

**Fig. 9.**
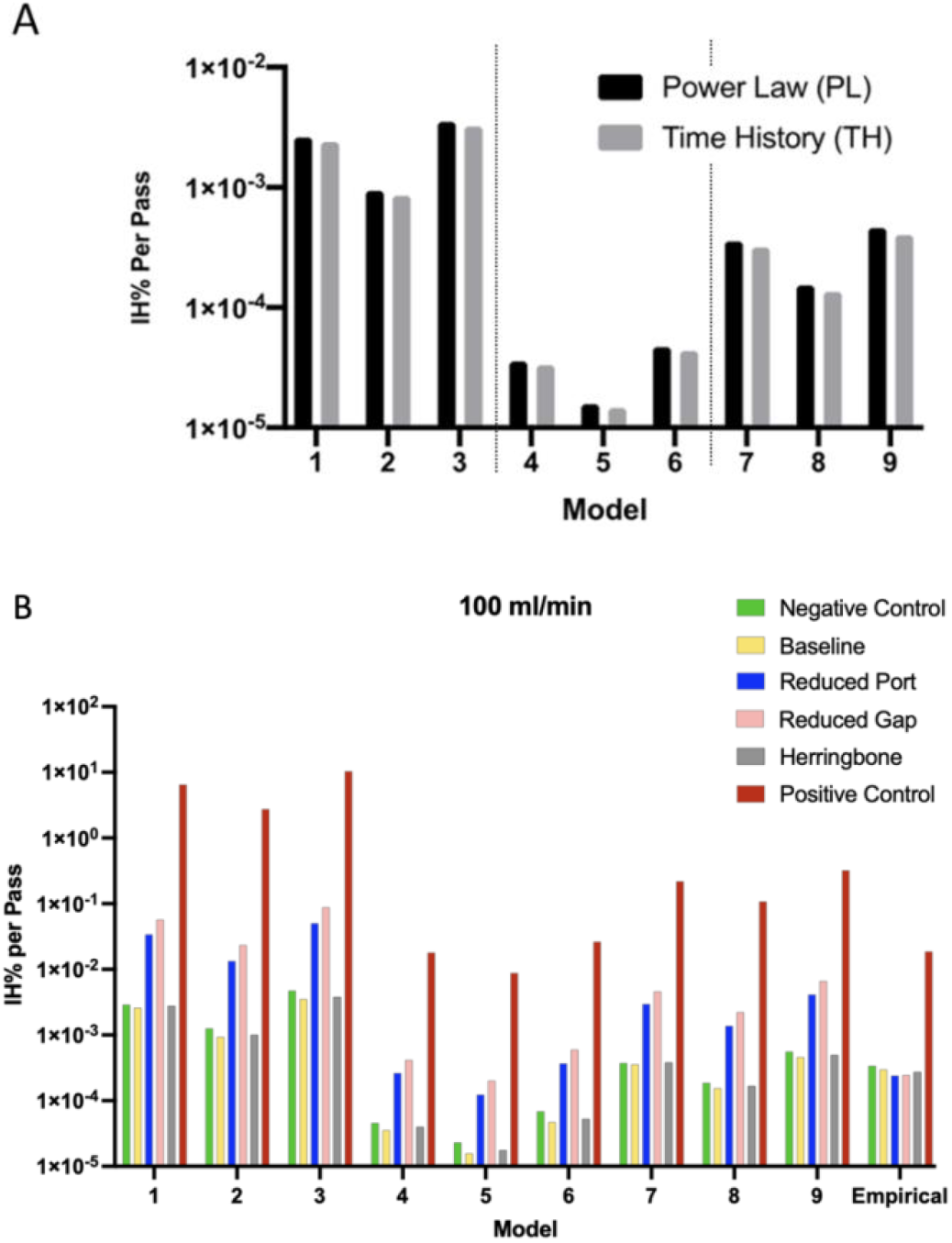
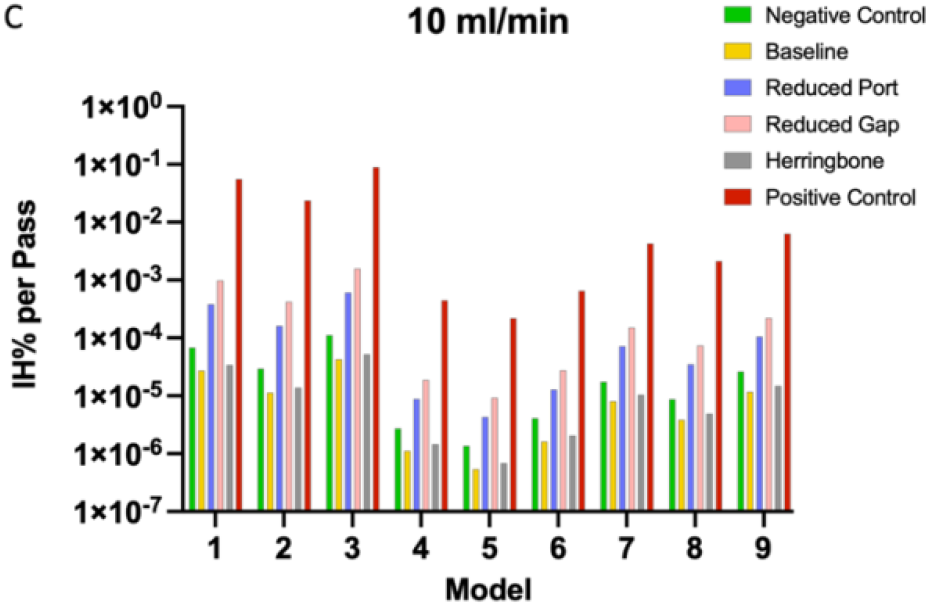
(A) Comparison of hemolysis data for the Baseline device for both PL and TH models (total 18 models) with Giersiepen parameters (Models 1-3), Heuser/Opitz parameters (Models 4-6), and the Zhang parameters (Models 7-9) (B, C) simulation hemolysis values for the Power-law models for all four devices and controls for flow rates 100 ml min^-1^ (B) and 10 ml min^-1^ (C) respectively. Panel B also includes the empirical hemolysis values.

### Empirical: Hemolysis

Fig 10A shows representative data from one day of empirical experiments. Hemolysis is nearly linear with the number of passes. It is evident that all devices exhibit significantly lower levels of hemolysis (<1.5 × 10^−4^ IHPP) compared to the positive control (∼1× 10^−2^IHPP). Fig. 10B presents the mean hemolysis values of all experiment days. It was observed that even without a device attached to the circuit (tubing only), the mean hemolysis values were found to be statistically comparable to the values that which the devices were attached (p > 0.05). This suggests the device-only hemolysis (excluding pump, tubing, and connectors), calculated as the difference between the maximum device-induced damage and the tubing-only circuit is 1.5 × 10^−4^ IHPP (3 × 10^−4^– 1.5 × 10^−4^ = 1.5 × 10^−4^ IHPP). As the empirical hemolysis values for 100 ml min^-1^ flow rates were so low, we determined that 10 ml min^-1^ flow conditions would generate hemolysis well below our measurement sensitivity, thus empirical experiments were not performed for 10 ml min^-1^. While comparing within the empirical cases, the baseline hemolysis value was not statistically distinguishable from the other devices (herringbone, reduced port, and reduced gap) (p > 0.05). The positive control IHPP values on the other hand was statistically significantly different from all other devices (p < 0.05). Table S1 reports the cross-device comparison p values obtained from the one-way ANOVA test.

**Fig. 10.**
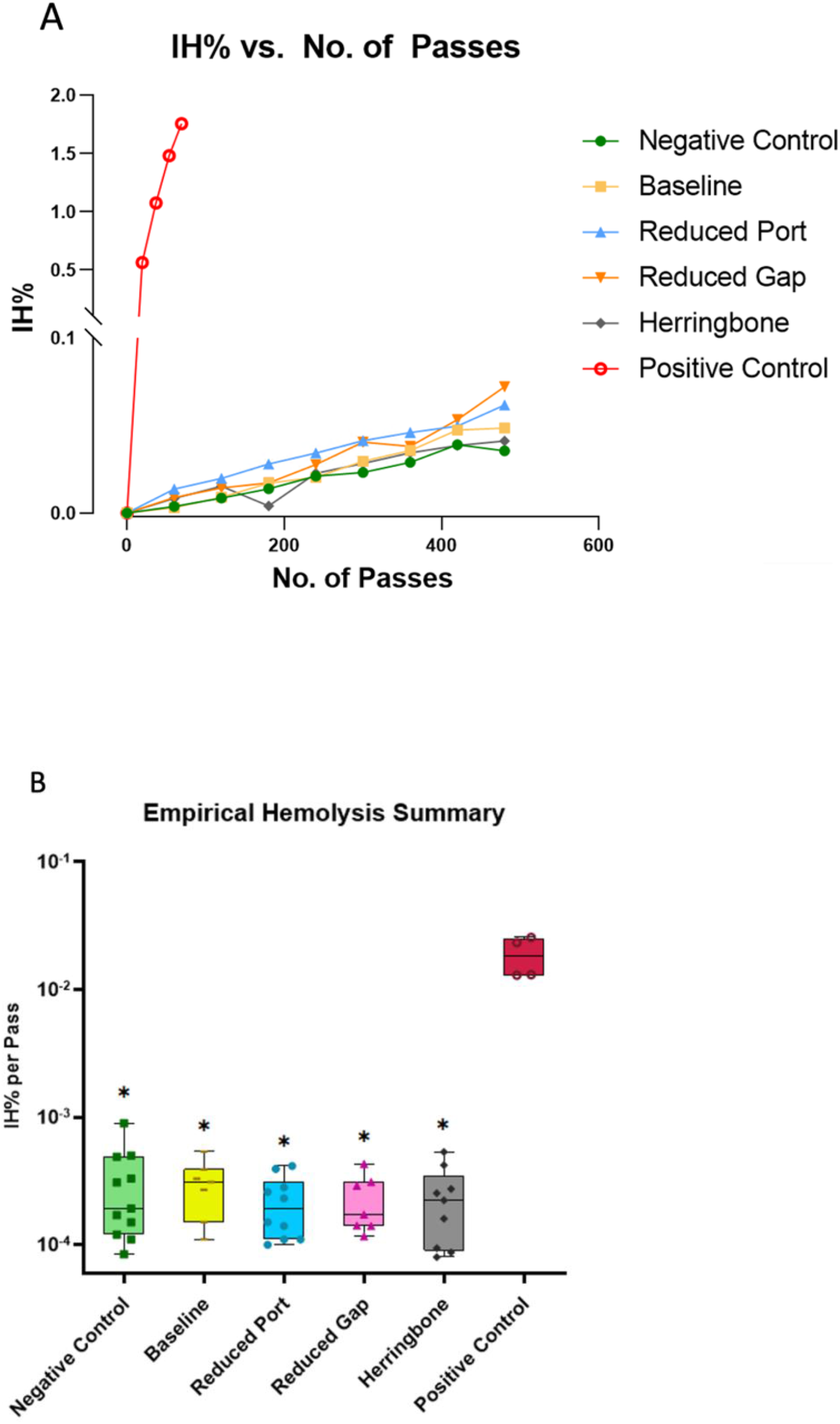
(A) IH% Vs. No. of Passes plot for one empirical experiment, (B) Overall mean of empirical data comparing IHPP for each device and controls. The solid midline represents the median IHPP and the whiskers are the lower and upper range of IHPP. Boxes containing asterisks depicts significantly different IHPP value from the positive control (p < 0.005).

**Fig. 11.**
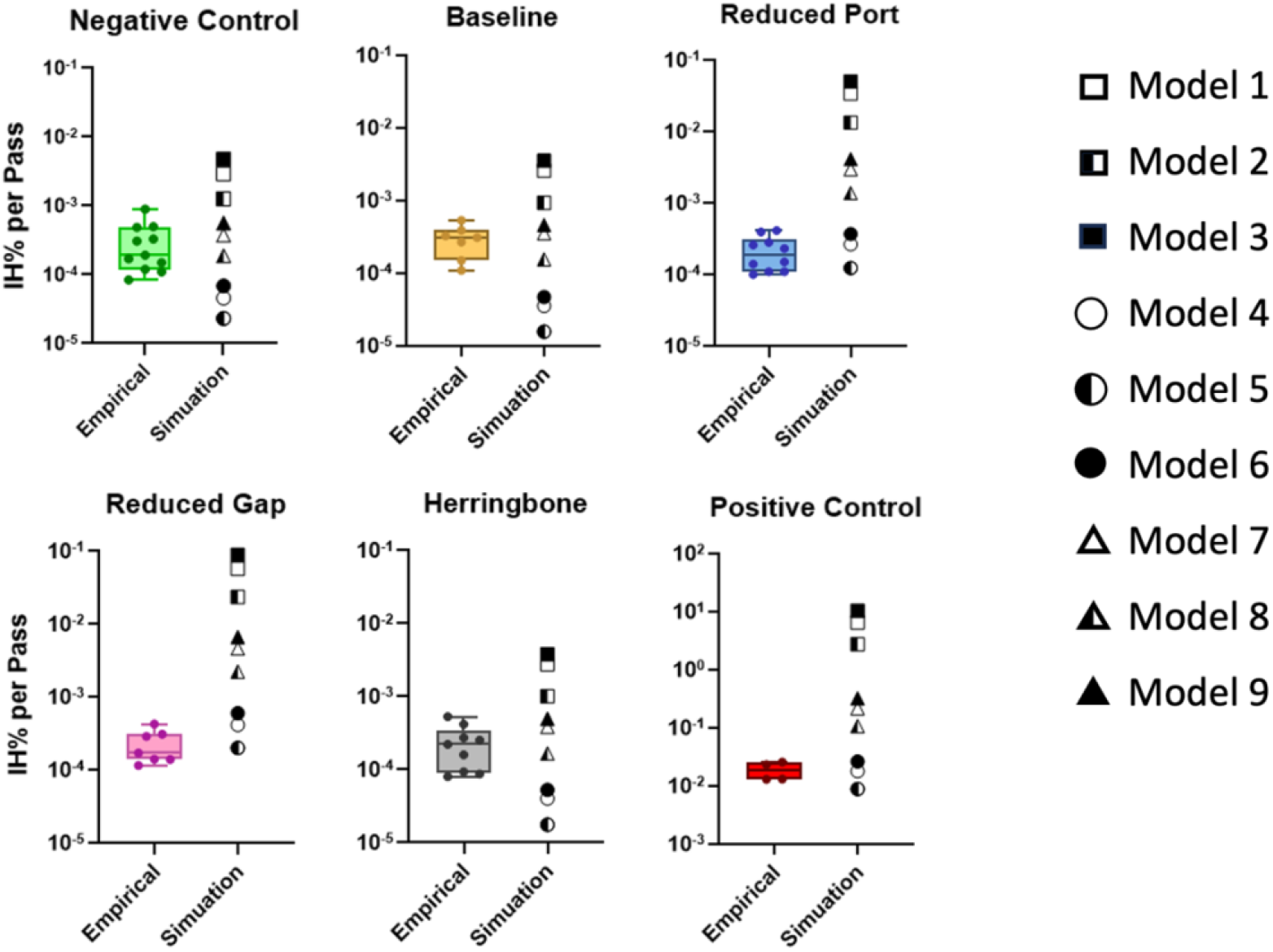
IH% per pass for individual devices and controls: Empirical vs. Simulation. The whiskers are the lower and upper range of IHPP. The squares, circles, and triangles represent Giersiepen (models 1 to 3), Heuser-Optiz (models 4 to 6), and Zhang (models 7 to 9) PL models respectively.

### Empirical vs. Simulation Hemolysis

For all cases (devices and controls) models (1, 2, 3) overpredicted (IHPP to IHPP) the hemolysis values when compared to the empirical (Fig. 8). Models (4, 5, 6) underpredicted (to IHPP) for the negative control, baseline, and herringbone device, but the prediction they provided for relatively higher shear devices: reduced port, reduced gap, and positive control, were rather accurate. Likewise, models (7 and 8) that best agreed for negative control, baseline, and herringbone device overpredicted (to 0.45 IHPP) for reduced port, reduced gap, and positive control. Model 9 followed a similar trend as models (7 and 8) with lesser accuracy. Table S2 reports the empirical vs. simulation comparison p values obtained from one sample t-test.

## 6. Discussion

The models predicted hemolysis values across several orders of magnitude, which is consistent with the literature.[26] It was observed that the time history model results in no significant improvement in the accuracy of prediction. This is expected because the time-history effect should be small since the magnitude of hemolysis predicted was low. Our observations revealed that for devices with average shear rates below 500 s^-1^ (baseline, herringbone, and negative control), models 7 and 8 exhibited the closest predictions. Conversely, for devices operating at shear rates equal to or exceeding 600 s^-1^ (reduced port, reduced gap, and positive control), models 4-6 demonstrated the closest predictions. This relationship suggests that when designing a microfluidic device expected to operate in a low shear range, selecting models 7 and 8 for hemolysis prediction is optimal. Conversely, when the prototype device is anticipated to operate in a high shear range, selecting models 4-6 is ideal. Although beyond the scope of this work, recent studies have presented methods to develop device-specific parameters using empirical data that spans multiple flow rates and a range of device geometries.[73] They determined device-specific model parameters, through a high number of empirical replicates and Kriging surrogate modeling, that produce hemolysis values that are closer to the empirical findings.[73]

The coefficients for the Giersiepen models (models 1-3) were derived from experiments encompassing shear rate ranges from 0 s^-1^ to 70833 s^-1^, for the Heuser/Optiz models (models 4-6) the shear rate ranged from 10000 s-1 to 166700 s-1, and for the Zhang models (models 7-9) the shear rate ranged from 13890 s-1 to 88890 s-1. Notably, the Heuser/Optiz models exhibited the highest shear rate ranges while calculating their coefficients, and as mentioned earlier, these models demonstrated good agreement with high-shear devices. This observation might suggest that the coefficients determined by Heuser/Optiz, being derived from shear rate ranges closest to those employed by high-shear devices, resulted in a closer agreement with these devices. Conversely, for the other models, such a clear correlation is not evident. This lack of correlation might be attributed to the differences in shear application methods utilized by those models compared to our approach of using microfluidics. Other factors, such as differing blood characteristics may also contribute to differences between empirical data and the previously derived models. In summary, the variations in the experimental shear rate ranges and the methods of shear application among different models can potentially impact their correlation with specific devices, especially in the context of microfluidics-based studies. A future work could be to recommend new sets of coefficients derived from microfluidic shear applications, which can lead to the development of more accurate hemolysis prediction models for microfluidics.

For all computational models, the three device modifications yielded an increase in hemolysis over the baseline device. This means the models were sensitive to each of the three design changes introduced and successfully predicted an increase in hemolysis for all three, with the largest incremental increase resulting from the two designs intended to increase shear (reduced port and reduced gap). The models may be useful for device design if multiple devices are required to be compared with each other in terms of their hemolytic tendency.

Empirical hemolysis observed for all four microfluidic devices was remarkably low (< 2 ppm hemolysis) and statistically indistinguishable for all four design iterations when compared to the negative control (p > 0.05) (Fig. 7). The low device-induced damage within all devices is promising for clinical use. A maximum of 3 ppm pass^-1^ of RBCs are damaged by the device, tubing and pump combined, which is lower than conventional ECMO loops (∼ 10 ppm pass^-1^). [74] The incremental damage caused by any of these devices is less than 1.5 ppm pass^-1^. Although these devices contain sharp edges and small features that induce non-physiological shear stresses at the tested flow rates, these stresses appear insufficient to induce significant hemolysis in the devices. The experimental conditions (flow rate: 100 ml min^-1^, reservoir volume: 60 mL, experiment length: 5-8 hours) generate over 650 passes through the devices, which is comparable to the time required for short-term ECMO support as a “bridge”. [75] The number of passes through the device in these experiments may greatly exceed what would normally be experienced if used clinically as a wearable continuous-flow device.

As previously stated, the results of empirical hemolysis for all four devices did not demonstrate a significant difference when compared to the negative control. This suggests that the hemolysis induced by the test loop and syringe pump may have overshadowed any hemolysis induced by the devices themselves. Therefore, the device-induced hemolysis level was deemed negligible for the tested devices. However, it is important to note that the positive control hemolysis was considerably higher, as confirmed by both empirical and computational analyses. Although the computational models did not accurately predict subtle differences in hemolysis between the devices, they were able to differentiate high levels of hemolysis induced by the positive control. This indicates that for microfluidic design iterations, the model can predict the trend of hemolysis accurately if the modeled hemolysis exceeds a certain threshold IHPP. Based on our experiments, this threshold is currently set at the hemolysis level obtained from the positive control. In future studies, it will be necessary to refine this threshold value and determine the lowest IHPP for computational analyses to align with empirical hemolysis measurements. Furthermore, future experiments should consider incorporating both computational and empirical testing of thrombosis for the devices to enhance hemocompatibility assessment.

The use of multi-functional or multi-physical models to guide device design is promising and increasingly practical with advancements in techniques and computer power. The ability to study design variants through simulation leads to less expensive and faster design iteration, eventually resulting in better-optimized designs. The low device-induced hemolysis (< 2 ppm) measured in our study at physiologically relevant flow rates is promising for the future development of microfluidic dialyzers and oxygenators, and our work demonstrates that computational modeling may supplement empirical testing to expedite the design optimization of these devices. Further, the study indicates that the power law has a higher tendency to over-predict hemolysis in microfluidic devices, and further customization is required to implement these existing models to a microfluidic device geometry.

## Acknowledgments

SiMPore Inc. for the design of baseline device variant. NIH award number 1R21HL156143-01 to SWD and NSF award number IIP 1660177 to TRG.

## Supplementary

**Table S1:**
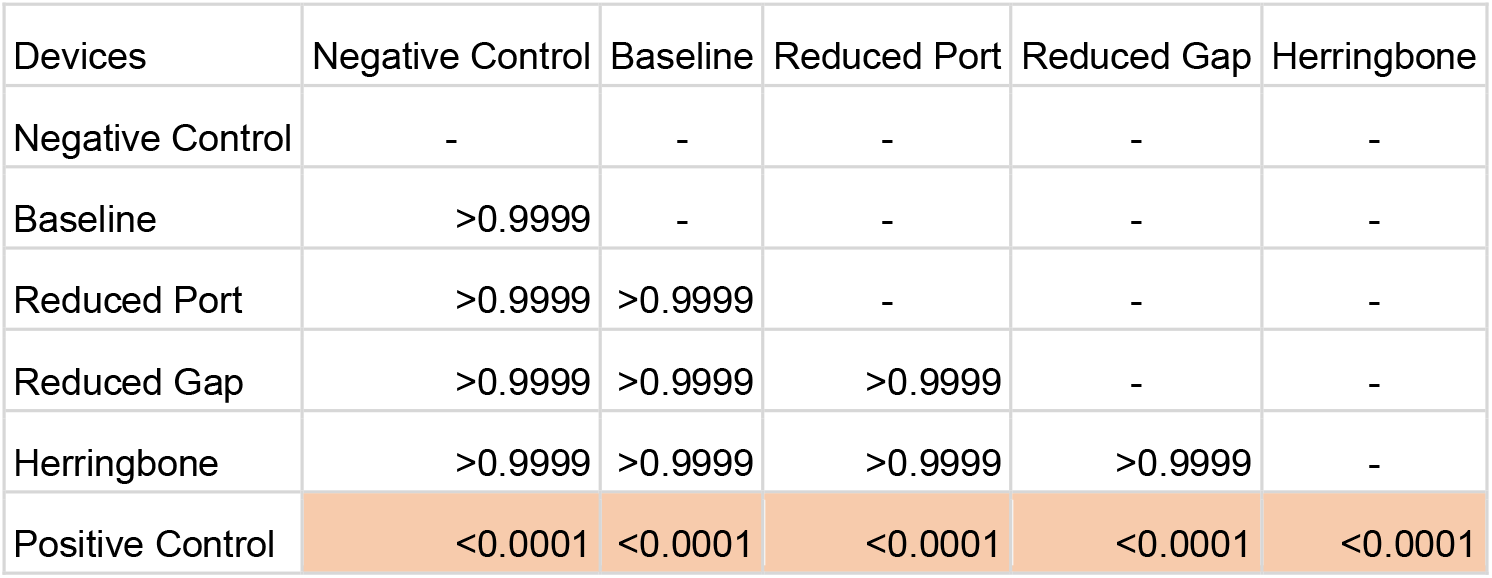
One-way ANOVA p values, (cross-device comparison of empirical values), boxes highlighted in orange (p < 0.05) means the IHPP difference was statistically significant. Whereas the non-highlighted boxes represent the differences not to be statistically significant.

**Table S2:** One sample t-test p values, (empirical vs model), boxes highlighted in green (p > 0.05) means the difference between the IHPP of empirical and model was not statistically significant. Whereas the non-highlighted boxes represent statistically significant differences.

|  | Models |  |  |  |  |  |  |  |  |
| --- | --- | --- | --- | --- | --- | --- | --- | --- | --- |
| Devices | 1 | 2 | 3 | 4 | 5 | 6 | 7 | 8 | 9 |
| Negative Control | <0.0001 | <0.0001 | <0.0001 | 0.0056 | 0.0034 | 0.0095 | 0.3538 | 0.1461 | 0.0056 |
| Baseline | <0.0001 | <0.0001 | <0.0001 | 0.0029 | 0.002 | 0.0036 | 0.3261 | 0.0384 | 0.0237 |
| Reduced Port | <0.0001 | <0.0001 | <0.0001 | 0.2413 | 0.0301 | 0.0028 | <0.0001 | <0.0001 | <0.0001 |
| Reduced Gap | <0.0001 | <0.0001 | <0.0001 | 0.0053 | 0.567 | 0.0001 | <0.0001 | <0.0001 | <0.0001 |
| Herringbone | <0.0001 | <0.0001 | <0.0001 | 0.0057 | 0.0031 | 0.0081 | 0.0216 | 0.2246 | 0.001 |
| Positive Control | <0.0001 | <0.0001 | <0.0001 | 0.8388 | 0.0592 | 0.106 | <0.0001 | 0.0001 | <0.0001 |

**Table S3:** Device shear rates for 100 ml/min flow rate.

| Device | Volm. Avg. Shear Rate (1/s) | Max. Shear Rate (1/s) |
| --- | --- | --- |
| Negative Control | 284 | 596 |
| Baseline | 363 | 37430 |
| Reduced Port | 597 | 150878 |
| Reduced Gap | 947 | 105511 |
| Herringbone | 486 | 51482 |
| Positive Control | 14267 | 273768 |

**Table S4:**
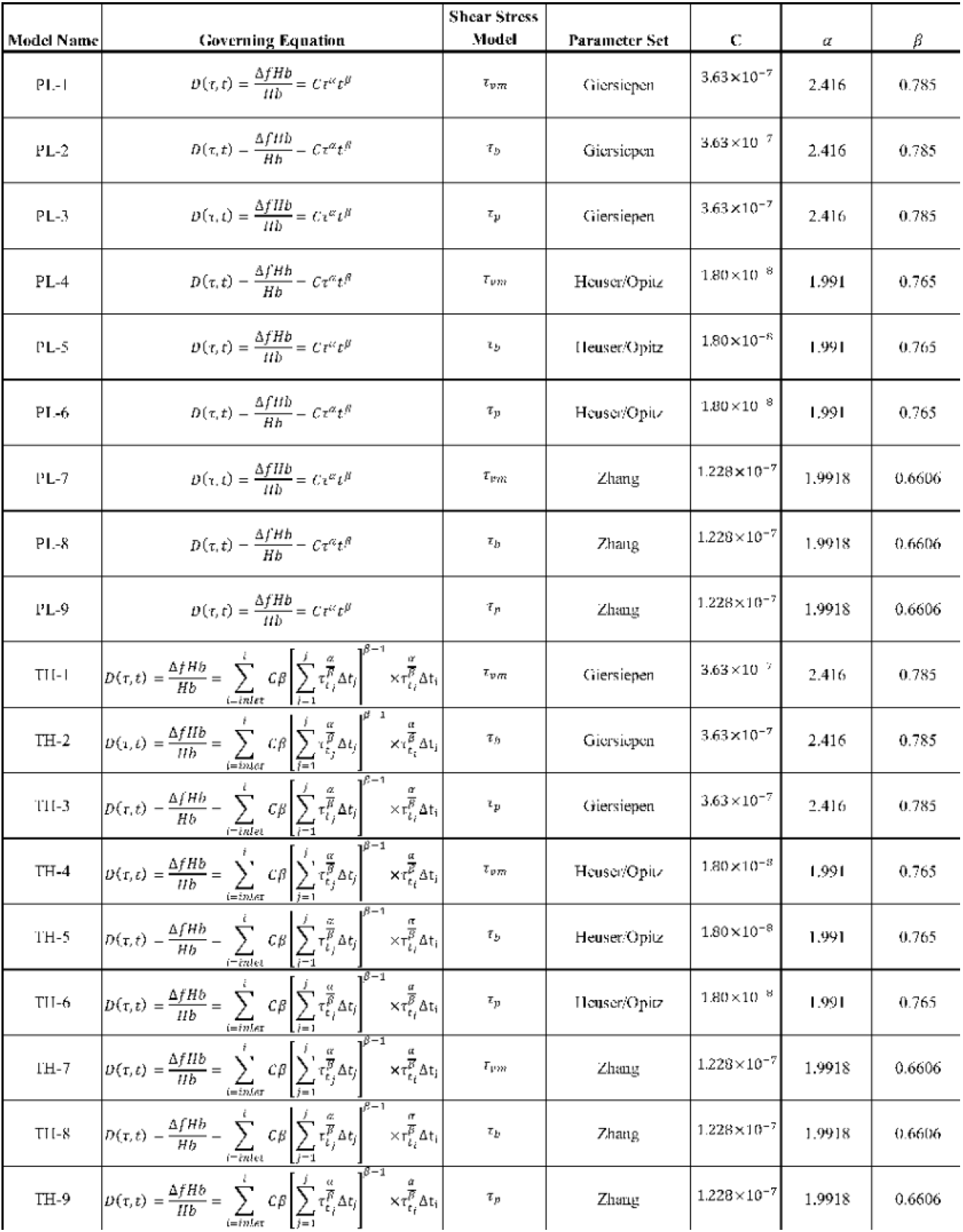
Properties of each hemolysis model tested *in-silico*.

